## Supplementary 1-6 for "Life-history adaptation under climate warming magnifies the agricultural footprint of a cosmopolitan insect pest"

**Index:**

**Supplementary 1:** *Predictions for how thermal adaptation via increased energy acquisition affects the agricultural footprint of insect pests.*

**Supplementary 2:** *Additional analyses of gene expression data*

**Supplementary 3:** *Univariate models of thermal adaptation in life history traits*

**Supplementary 4:** *Estimates of laboratory fitness in experimental evolution lines*

**Supplementary 5:** *Estimates of host consumption in experimental evolution lines*

**Supplementary 6**: *Selection on female body mass at different temperatures*

*.*

**Supplementary 1:**

*Predictions of how thermal adaptation via increased resource acquisition affects the agricultural impact of insect pests based on protein biophysics.*

Biological rates of ectotherms show an empirically well-described relationship with temperature that closely mirrors the thermodynamic performance of enzymes (Hochachka and Somero 2002; Angilletta 2009), because biological rates are governed at the biochemical level by the enzymatic reaction rate, $k_{cat}$ (Evans and Polanyi 1935; Eyring 1935):

$k_{cat}\propto A_{B}e^{-Ea/RT}$ $r=r_{0}e^{-\Delta G^{a}/RT}$	(Eq. S1)

where *E_a_* is the activation energy required for the enzymatic reaction to occur, $R$ is the universal gas constant (0.002 kcal mol^-1^), *T* is temperature in Kelvin and *A*_B_ is a species and rate specific constant for enzyme catalys (Clarke 2004; Dell et al. 2011). Equation 1 thus describes an increase in reaction rate with temperature (**Fig. 1a**, main text). The observed decline in biological rate at temperatures exceeding the organism’s thermal optimum is attributed to a reduction in the proportion of functional enzyme due to reversable inactivation via protein unfolding (Bloom et al. 2005; DePristo et al. 2005; Bershtein et al. 2017; Echave and Wilke 2017; Agozzino and Dill 2018) (**Fig. 1a**). The proportion of folded protein ready to do work in a biological process follows a Boltzmann probability as a function of the Gibbs free energy of folding, $\Delta G$, which is thus a measure of protein stability (DePristo et al. 2005):

P_fold_ = $\prod_{i=1}^{n} \frac{1}{\left. 1+A_{D,i}e^{{{\Delta G}_{i}(T)}/{RT}} \right.}$.	(Eq. S2)

Where constant *A*_D_ and stability, $\Delta G$, is given for each of a set of *n* proteins that act in sequence and contribute multiplicatively to a biological rate (Chen and Shakhnovich 2010; Echave and Wilke 2017). At a benign temperature of 25˚C, most natural proteins occur in functional (properly folded) state and the mean value of the Gibbs energy is negative (mean $\Delta G$ ~ -7 kcal mol^-1^; (DePristo et al. 2005; Chen and Shakhnovich 2009). However, $\Delta G$ is comprised of an enthalpy term ($\Delta H$) and a temperature-dependent entropy term ($\Delta S$) (Eyring and Polanyi 2013) that reduces to: $\Delta G\left( T \right)=\Delta H-T\Delta S$ over the ecologically relevant temperature range of most organisms (Chen and Shakhnovich 2010). Based on values from the literature (Chen and Shakhnovich 2010; Dill et al. 2011), *∆S* $\Delta\Delta S^{f}$~ $-0.25$ to $-$0.50 kcal/mol K^-1^ for a protein of typical length. From equation S2 it is therefore clear that warm temperatures increase entropy making $\Delta G$ less negative, leading to a rapid non-linear reduction in the fraction of functional protein (**Fig. 1a**).

The reaction rate kinetics of equation S1 have been combined with the protein folding stability of equation S2 to describe temperature-dependent biological rates and even fitness in simple scenarios (Bershtein et al. 2017; Echave and Wilke 2017). We here place these biophysical predictions into a life history framework incorporating acquisition and allocation trade-offs (Jong and Noordwijk 1992) to predict the temperature dependence of population growth rate:

$r(T)\propto$ ${{(1-c)}^{M}*M}^{b}*k_{cat}*$P_fold_	(Eq. S3)

where *M* is accumulated energy reserves, *c* is the survival cost of acquiring energy, and *b* is the allometric exponent for how growth and reproduction scale with *M*.

While there is variation in biological rates across organisms and for different traits (Clarke 2004; Angilletta 2009; Huey and Kingsolver 2011), inherent enzyme properties governing the temperature-dependence of protein stability and reaction rate have been shown to evolve slowly (Dill et al. 2011). Instead, thermal adaptation often proceeds via costly compensatory responses involving upregulation of molecular chaperones such as heat-shock proteins that aid protein folding at hot temperature (Feder et al. 2000), and enzyme catalysts that allow reactions to take place at cold temperature despite thermodynamic constraints (Hochachka and Somero 2002). Here we illustrate how such thermal compensation via increased energy acquisition can affect growth rates of ectotherm pests facing changing climates (see **Fig. 1C** in main text). Assume that the energy reserves, *M*, of an organism can be allocated with fraction *p* to produce molecules that increase enzyme reaction rate, and with fraction *q* to molecules that aid protein folding stability, leaving *M*(1-*p*-*q*) reserves for growth and reproduction. Inserting these expressions into equation S3 gives:

${r^{'}\left( T \right)= {(1-c)}^{M}*[M(1-p-q)]}^{b}*pMA_{B} e^{-\frac{Ea}{RT}}*\prod_{1}^{i} \frac{1}{\left. 1+\frac{A_{D}}{qM}e^{{{\Delta G}_{i}(T)}/{RT}} \right.}$	(Eq. S4)

We can find the optimal strategy of energy acquisition (increasing *M* at cost *c*) and allocation (of *M* to *p* and *q*) at different temperatures and for different costs of growth by numerically solving for the combination of *M*, *p* and *q* that maximize $r^{'}$.

Eq. S4 is well-supported in its qualitative sense. Eqs. S1 and S2 have received strong empirical support from in vitro measures of enzymes (Dill et al. 2011; Echave and Wilke 2017; Agozzino and Dill 2018) and recapitulates the asymmetry of empirically estimated thermal reaction norms of quantitative traits (Angilletta 2009). Thermal compensation via molecular chaperones is also well-understood (Hochachka and Somero 2002) and has been shown to trade-off with growth and reproduction (Feder et al. 2000). However, the quantitative solution to the multidimensional trade-off described by eq. S4 depends on the mechanistic basis of thermal sensitivity down to its finer details and exactly how energy reserves invested in thermal compensation affect folding stability and reaction rate, relationships that are not empirically well-defined nor completely understood (e.g. Eqs. S1 and S2 describe phenomenological rather than mechanistic relationships). Thus, our aim is only to provide qualitative predictions. We do this by using eq. S4 to illustrate how thermal niches can evolve via compensatory energy acquisition and allocation under a relatively high (*c* = 0.9) or low (*c* = 0.8) cost of energy acquisition, when an organism is adapting to a cold (23°C) or hot (35°C) temperature (**Fig. 1C**, main text). For other parameters we first assumed values that correspond to those commonly observed in the literature: ΔG_T=298K_ = -7 kcal mol^-1^; *∆S* $\Delta\Delta S^{f}$~ $-0.25$ kcal mol^-1^ K^-1^; *n* = 100 proteins; *b* = 0.75, *Ea* = 20 kcal mol^-1^(~ 0.7eV). *A_B_*, and *A_D_* were set to 1 for simplicity.

After finding the optimal strategy in the four scenarios, resulting thermal reaction norms for population growth rate were calculated and are depicted in **Fig. 1C** of the main text, along with the optimal energy acquisition strategy in each scenario. Cold-adapted populations outcompete hot-adapted population at cold temperature, but growth rates and the predicted impact on agriculture are greater at warm temperatures and for the hot-adapted population due to reduced thermodynamic constraints on reaction rates (Gillooly et al. 2002; Brown et al. 2004). Lower costs of feeding favour increased energy acquisition and allocation to maintenance, which improves thermal performance and increases thermal niche breadth (**Fig. 1C**). Increased feeding effort is particularly beneficial at hot temperature because there are increased demands on maintaining protein folding at these temperatures (**Fig. 1A**). Thermal adaptation via increased energy acquisition mediates an even greater impact on the rate of host consumption (given by the product of $r'$ and *M –* the latter assumed to be proportional to per capita host consumption) (**Fig. 1D, Fig. S1C-F**).

To further illustrate how warming temperatures affect energy acquisition and the predicted impact on agriculture, we calculated the optimal life-history strategy (*M’, p’* and *q’*) at each temperature under different assumptions of the thermal sensitivity of protein folding (**Figure S1**, *below*). We then calculated the associated population growth rate and rate of host consumption for each temperature and scenario. This confirmed that, if increased energy acquisition and allocation can compensate effects of temperature on molecular failure rates, greater thermal sensitivity favours increased foraging effort **(Fig. S1A, B**). Increased thermal sensitivity implies low fitness at high temperature per definition, so population growth rates remain low in this scenario (**Fig. S1C, D**) as thermal compensation via increased growth effort also comes at a cost (*c*). However, if costs of feeding are low, increased energy acquisition is more favourable, which increases population growth rate, and even more so, agricultural impact (**Fig. S1E, F**). In fact, the strategy to increase *M* at stressful temperature causes populations experiencing temperatures slightly hotter than those that maximize population growth rate to have the greatest agricultural impact (*compare* **Fig. S1C** vs. **E** & **Fig. S1D** vs. **F**).

We note that protein folding can also be compromised by acute cold stress (Feder et al. 2000; Hochachka and Somero 2002). However, such cold temperatures are typically not experienced during the active seasons of tropical and sub-tropical insects (Deutsch et al. 2008; Johansson et al. 2020), as the case for *C. maculatus* (Baur et al. 2023), and were therefore not modelled. Likewise, stressful temperatures affect organisms in a multitude of ways. Thus, our particular example using protein folding is merely meant to illustrate a general principle and was chosen based on the simple fact that the modelled relationships are empirically well-supported in qualitative sense (see above). Other physiological phenomena that scale with temperature, such as metabolic expenditure (Gillooly et al. 2001) and the production of harmful reactive oxygen species (Dowling and Simmons 2009), are likely to simultaneously contribute to increasing molecular failure rates at stressful temperature.

Finally, similar to previous models (Bershtein et al. 2017; Echave and Wilke 2017), we here assumed that growth and reproduction of complex organisms are proportional to the rate at which they generate functional enzyme, which is clearly simplistic considering the ecological complexity affecting the link between rates of biochemical reactions and population growth rate. Yet, this simplicity allows forming general predictions and is supported qualitatively by the fact that a range of different biological rates scale predictably with temperature (Brown et al. 2004; Angilletta 2009). All R code used to produce predictions is available as Supplementary

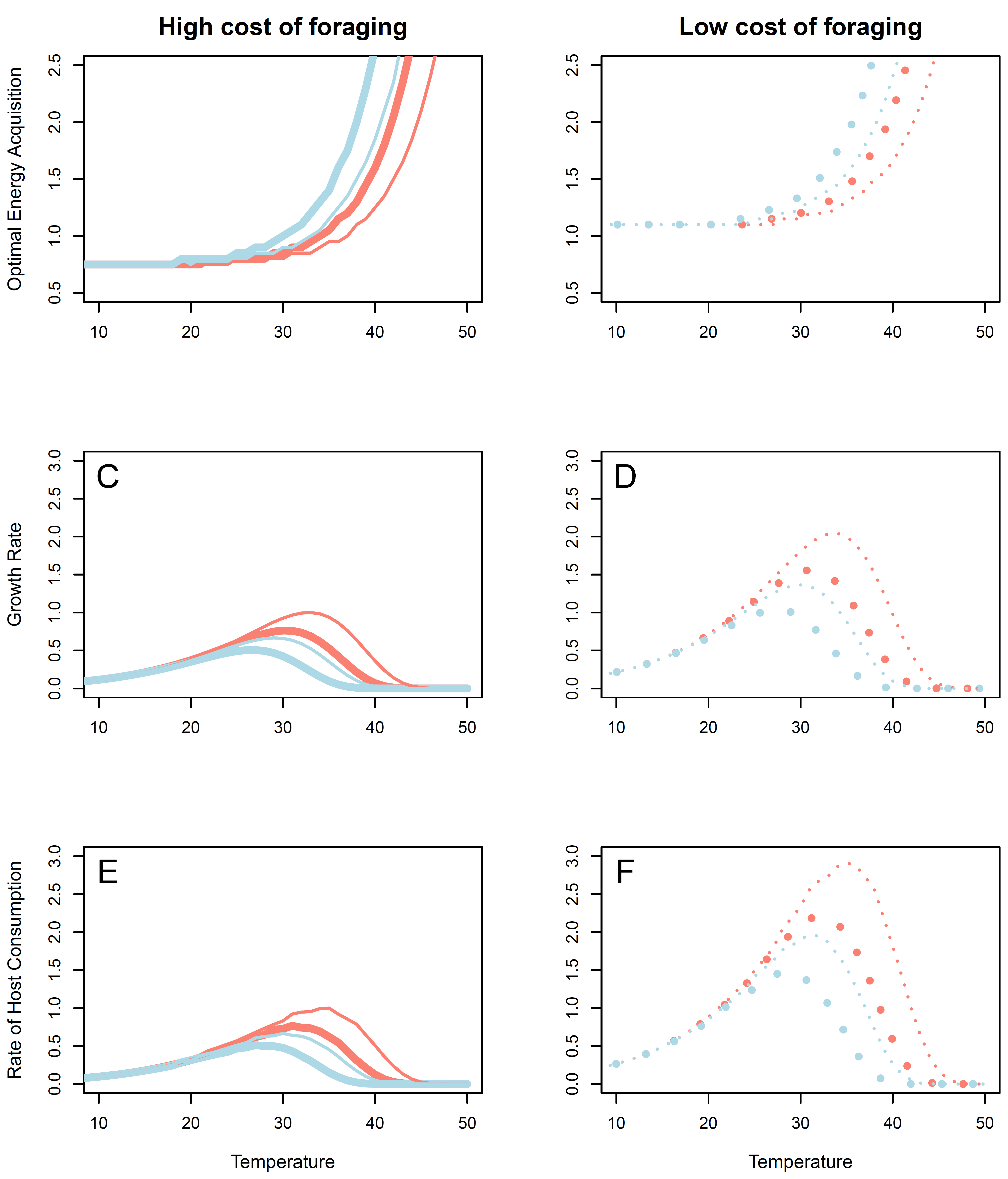

**Figure S1: Optimal energy acquisition and its agricultural impact**

In **A)** and **B),** the optimal strategy of energy acquisition (*M’*) at each temperature in a scenario of high (*c* = 0.9; left panels) or low (*c* = 0.8; right panels) costs of foraging. 100 (thin lines) or 300 (thick lines) multiplicatively acting proteins were assumed to contribute to growth and/or reproduction. These proteins were either relatively stable (ΔG_T=298K_ = -7 kcal mol^-1^; salmon) or less stable (ΔG_T=298K_ = -6 kcal mol^-1^; light blue). More numerous, and less stable, proteins contribute to increased thermal sensitivity and benefit increases in *M’.* In **C)** and **D),** the resulting population growth rate, and in **E)** and **F)** the rate of host consumption, are shown at each temperature for each scenario. Note that these results represent optimal solutions to the multidimensional life-history trade-offs and could either stem from genetic adaptation or adaptive plasticity. Results in panels C-F are scaled by the maximum of the scenario depicted by thin and full lines in salmon colour (ΔG_T=298K_ = -7 kcal mol^-1^; *n* = 100, *c* = 0.9).
**References:**

Agozzino, L., and K. A. Dill. 2018. Protein evolution speed depends on its stability and abundance and on chaperone concentrations. PNAS 115:9092–9097.

Angilletta, M. J. 2009. Thermal Adaptation: A Theoretical and Empirical Synthesis. OUP Oxford.

Baur, J., M. Zwoinska, M. Koppik, R. R. Snook, and D. Berger. 2023. Heat stress reveals a fertility debt owing to postcopulatory sexual selection. Evolution Letters qrad007.

Bershtein, S., A. W. Serohijos, and E. I. Shakhnovich. 2017. Bridging the physical scales in evolutionary biology: from protein sequence space to fitness of organisms and populations. Curr. Opin. Struct. Biol. 42:31–40.

Bloom, J. D., J. J. Silberg, C. O. Wilke, D. A. Drummond, C. Adami, and F. H. Arnold. 2005. Thermodynamic prediction of protein neutrality. Proceedings of the National Academy of Sciences of the United States of America 102:606–611.

Brown, J. H., J. F. Gillooly, A. P. Allen, V. M. Savage, and G. B. West. 2004. Toward a Metabolic Theory of Ecology. Ecology 85:1771–1789.

Chen, P., and E. I. Shakhnovich. 2009. Lethal Mutagenesis in Viruses and Bacteria. Genetics 183:639–650.

Chen, P., and E. I. Shakhnovich. 2010. Thermal adaptation of viruses and bacteria. Biophysical journal 98:1109–1118.

Clarke, A. 2004. Is there a Universal Temperature Dependence of metabolism? Functional Ecology 18:252–256.

Dell, A. I., S. Pawar, and V. M. Savage. 2011. Systematic variation in the temperature dependence of physiological and ecological traits. Proceedings of the National Academy of Sciences 108:10591–10596.

DePristo, M. A., D. M. Weinreich, and D. L. Hartl. 2005. Missense meanderings in sequence space: a biophysical view of protein evolution. Nature Reviews Genetics 6:678–687.

Deutsch, C. A., J. J. Tewksbury, R. B. Huey, K. S. Sheldon, C. K. Ghalambor, D. C. Haak, and P. R. Martin. 2008. Impacts of climate warming on terrestrial ectotherms across latitude. PNAS 105:6668–6672.

Dill, K. A., K. Ghosh, and J. D. Schmit. 2011. Physical limits of cells and proteomes. PNAS 108:17876–17882.

Dowling, D. K., and L. W. Simmons. 2009. Reactive oxygen species as universal constraints in life-history evolution. Proceedings of the Royal Society B: Biological Sciences 276:1737–1745.

Echave, J., and C. O. Wilke. 2017. Biophysical models of protein evolution: understanding the patterns of evolutionary sequence divergence. Annual review of biophysics 46:85–103.

Evans, M. G., and M. Polanyi. 1935. Some applications of the transition state method to the calculation of reaction velocities, especially in solution. Trans. Faraday Soc. 31:875–894. The Royal Society of Chemistry.

Eyring, H. 1935. The Activated Complex in Chemical Reactions. J. Chem. Phys. 3:107–115. American Institute of Physics.

Eyring, H., and M. Polanyi. 2013. On Simple Gas Reactions. Zeitschrift für Physikalische Chemie 227:1221–1246. De Gruyter.

Feder, M. E., A. F. Bennett, and R. B. Huey. 2000. EVOLUTIONARY PHYSIOLOGY1. 29.

Gillooly, J. F., J. H. Brown, G. B. West, V. M. Savage, and E. L. Charnov. 2001. Effects of Size and Temperature on Metabolic Rate. Science 293:2248–2251.

Gillooly, J. F., E. L. Charnov, G. B. West, V. M. Savage, and J. H. Brown. 2002. Effects of size and temperature on developmental time. Nature 417:70–73.

Hochachka, P. W., and G. N. Somero. 2002. Biochemical Adaptation: Mechanism and Process in Physiological Evolution. Oxford University Press.

Huey, R. B., and J. G. Kingsolver. 2011. Variation in universal temperature dependence of biological rates. Proceedings of the National Academy of Sciences 108:10377–10378.

Johansson, F., G. Orizaola, and V. Nilsson-Örtman. 2020. Temperate insects with narrow seasonal activity periods can be as vulnerable to climate change as tropical insect species. Sci Rep 10:8822. Nature Publishing Group.

Jong, G. de, and A. J. van Noordwijk. 1992. Acquisition and Allocation of Resources: Genetic (CO) Variances, Selection, and Life Histories. The American Naturalist 139:749–770.

**Supplementary 2:** Gene expression analyses

```
Supplementary Table 2a: Gene annotations for all heat stress and reproduction genes (see separate file).

Supplementary Table 2b: Gene ontology for all heat stress and reproduction genes (see separate file).


Supplementary Table 2c: Nested ANOVA of allocation response based on the 134 antagonistic genes (corresponds to Figure 3D of main text. The model was fit using the “aov” function and was specified as:

Allocation ~ Evolution.regime*temperature*Origin + Error(Population)

Error: Population
                         	Df    Sum Sq   Mean Sq F value   Pr(>F)
Regime             		2    1320027    660013   0.122 0.8867
Temperature                	2  140331406  70165703  12.972 0.0030 **
Origin                	2  311216068 155608034  28.768 0.0002 ***
Regime:Origin  		4  238677450  59669362  11.031 0.0024 **
Residuals                 	8   43272241   5409030

Error: Within
                     		Df    Sum Sq   Mean Sq F value   Pr(>F)
Temperature                     	2   4.3e+09  2.2e+09 262.982 1.63e-13 ***
Regime:Temperature             	4   1.7e+07  4.2e+06   0.514   0.7264
Temeprature:Origin              	4   9.5e+07  2.4e+07   2.867   0.0553 .
Regime:Temperature:Origin  	8   1.1e+08  1.4e+07   1.738   0.1610
Residuals                       	17  1.4e+08  8.3e+06
```

```
Supplementary Table 2d: Nested ANOVA of allocation response based on all the 21 genes upregulated in response to reproduction. The model was fit using the “aov” function and was specified as:

Allocation ~ Evolution.regime*temperature*Origin + Error(Population)

Error: Population
                          	Df Sum Sq Mean Sq F value   Pr(>F)
Regime                   	2 251496  125748  11.964 0.00394 **
Temperature               	2  23331   11666   1.110 0.37548
Origin                    	2  56967   28483   2.710 0.12629
Regime:Origin             	4 115818   28954   2.755 0.10385
Residuals                 	8  84084   10510

Error: Within
                     		Df    Sum Sq   Mean Sq F value   Pr(>F)
Temperature                      2 1117957  558978  72.545 4.74e-09 ***
Regime:Temperature               4   33218    8304   1.078   0.3983
Temperature:Origin               4   73571   18393   2.387   0.0918 .
Regime:Temperature:Origin        8   38745    4843   0.629   0.7433
Residuals                       17  130990    7705


Supplementary Table 2e: Nested ANOVA of allocation response based on all the 113 genes upregulated in response to heat shock. The model was fit using the “aov” function and was specified as:

Allocation ~ Evolution.regime*temperature*Origin + Error(Population)

Error: Population
                       Df Sum Sq Mean Sq F value   Pr(>F)
Regime                  2   2697138   1348569   0.251 0.783913
Temperature             2 137047700  68523850  12.755 0.003248 **
Origin                  2 305102729 152551364  28.396 0.000232 ***
Regime:Origin           4 235845545  58961386  10.975 0.002475 **
Residuals               8  42979006   5372376

Error: Within
                     		Df    Sum Sq   Mean Sq F value   Pr(>F)
Temperature                     	2   4.21e+09  2.1e+09  258.02 1.91e-13 *** Regime:Temperature             	4   1.63e+07  4.08e+06   0.500   0.736
Temperature:Origin              	4   9.14e+07  2.28e+07   2.800   0.059 .
Regime:Temperature:Origin  	8   1.15e+08  1.44e+07   1.761   0.155
Residuals                      	17  1.39e+08  8.16e+06
```

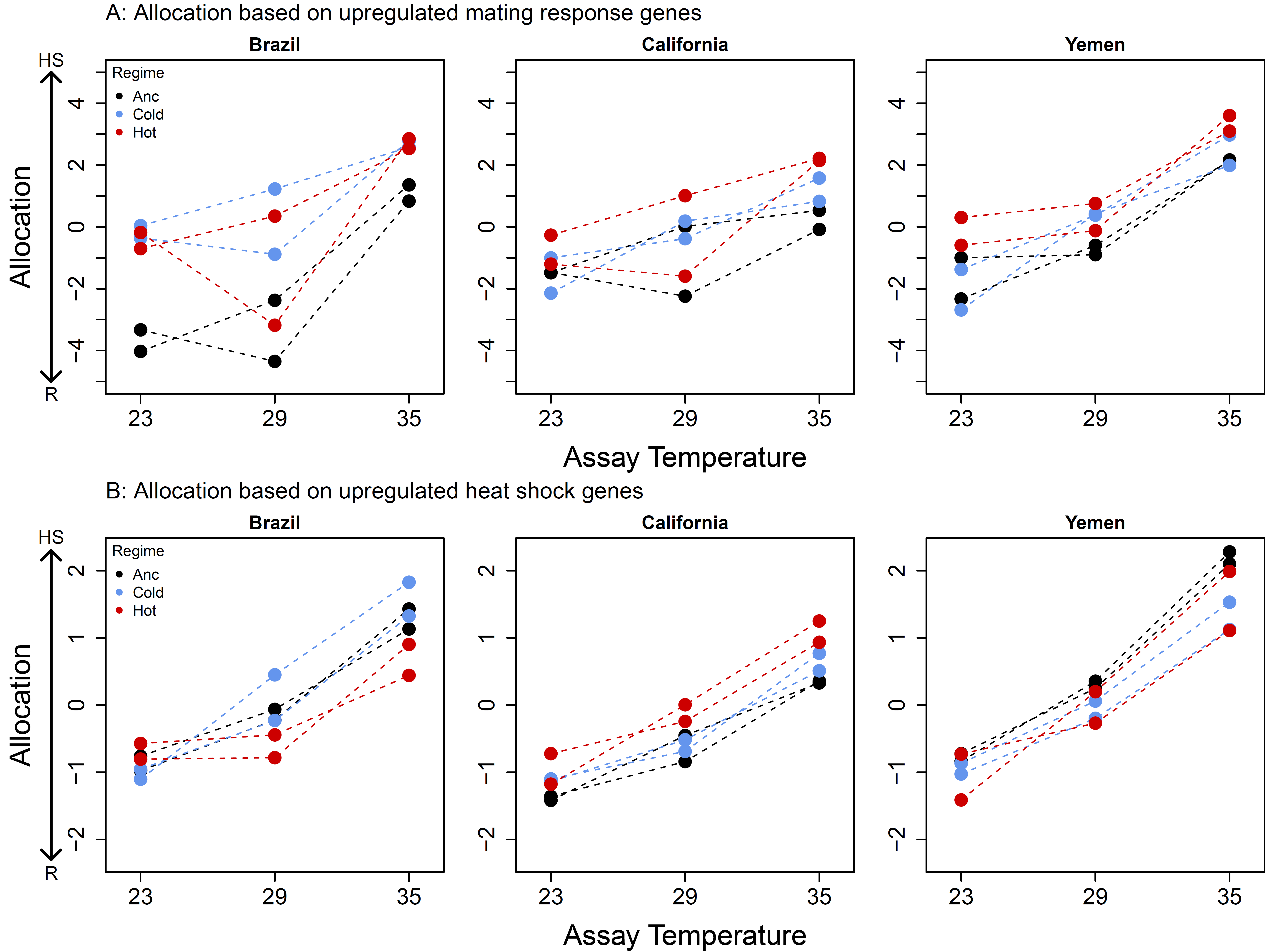

**Figure S2: Allocation trade-off between reproduction and heat stress response in experimental evolution lines based on two sets of overlapping genes.**

A: allocation based on expression of the 21 genes upregulated in response to mating in hot-adapted (red), cold-adapted (blue) and ancestors (black), reared at 23, 29, or 35°C. B: the same patterns for 113 genes upregulated in response to heat shock. Allocation towards stress response increased with assay temperature for both sets of genes. There was not general effect of evolution regime across the three backgrounds when analyzing the 113 genes upregulated in response to heat shock, while ancestors showed significantly higher expression of the 21 genes upregulated in response to mating.

```
Supplementary Table 2f: Nested ANOVA of allocation to reproduction based on all the 1269 differentially expressed genes in response to mating. The model was fit using the “aov” function and was specified as:

Allocation ~ Evolution.regime*Temperature*Origin + Error(Population)

Error: Population
                         	Df    Sum Sq   Mean Sq F value   Pr(>F)
Regime             		2    1.4e+10   6.8e+09   3.82  0.0685 .
Temperature                	2    9.1e+09   4.5e+09   2.52  0.142
Origin                	2    1.4e+10   6.8e+09   3.76  0.070 .
Regime:Origin  		4    6.2e+09   1.5e+09   0.86  0.527
Residuals                 	8    1.4e+10   1.8e+09

Error: Within
                     		Df    Sum Sq   Mean Sq F value   Pr(>F)
Temperature                     	2   7.9e+10  3.6e+10   23.11  1.4e-05 ***
Regime:Temperature             	4   4.8e+09  1.2e+09    0.69    0.61
Temeprature:Origin              	4   4.2e+09  1.1e+09    0.62    0.65
Regime:Temperature:Origin  	8   7.8e+09  9.7e+08    0.57    0.79
Residuals                      	17  2.9e+10  1.7e+09


Supplementary Table 2g: Nested ANOVA of allocation to stress response based on all the 765 differentially expressed genes in response to heat shock. The model was fit using the “aov” function and was specified as:

Allocation ~ Evolution.regime*temperature*Origin + Error(Population)

Error: Population
                         	Df    Sum Sq   Mean Sq F value   Pr(>F)
Regime             		2    5563026   2781513    1.31 0.32080  Temperature                	2   24185478  12092739    5.72 0.02872 * Origin                	2   93953143  46976571   22.21 0.00054 *** Regime:Origin  		4   50331286  12582822    5.95 0.01601 *
Residuals                 	8    16921967  2115246

Error: Within
                     		Df    Sum Sq   Mean Sq F value   Pr(>F)
Temperature                     	2   1.08e+09 5.42e+08  196.21 1.8e-12 *** Regime:Temperature             	4   8.28e+06 2.07e+06    0.75   0.572
Temeprature:Origin              	4   3.24e+07 8.10e+06    2.94   0.052 .
Regime:Temperature:Origin  	8   3.66e+07 4.58e+06    1.66   0.181
Residuals                      	17  4.69e+07 2.76e+06
```

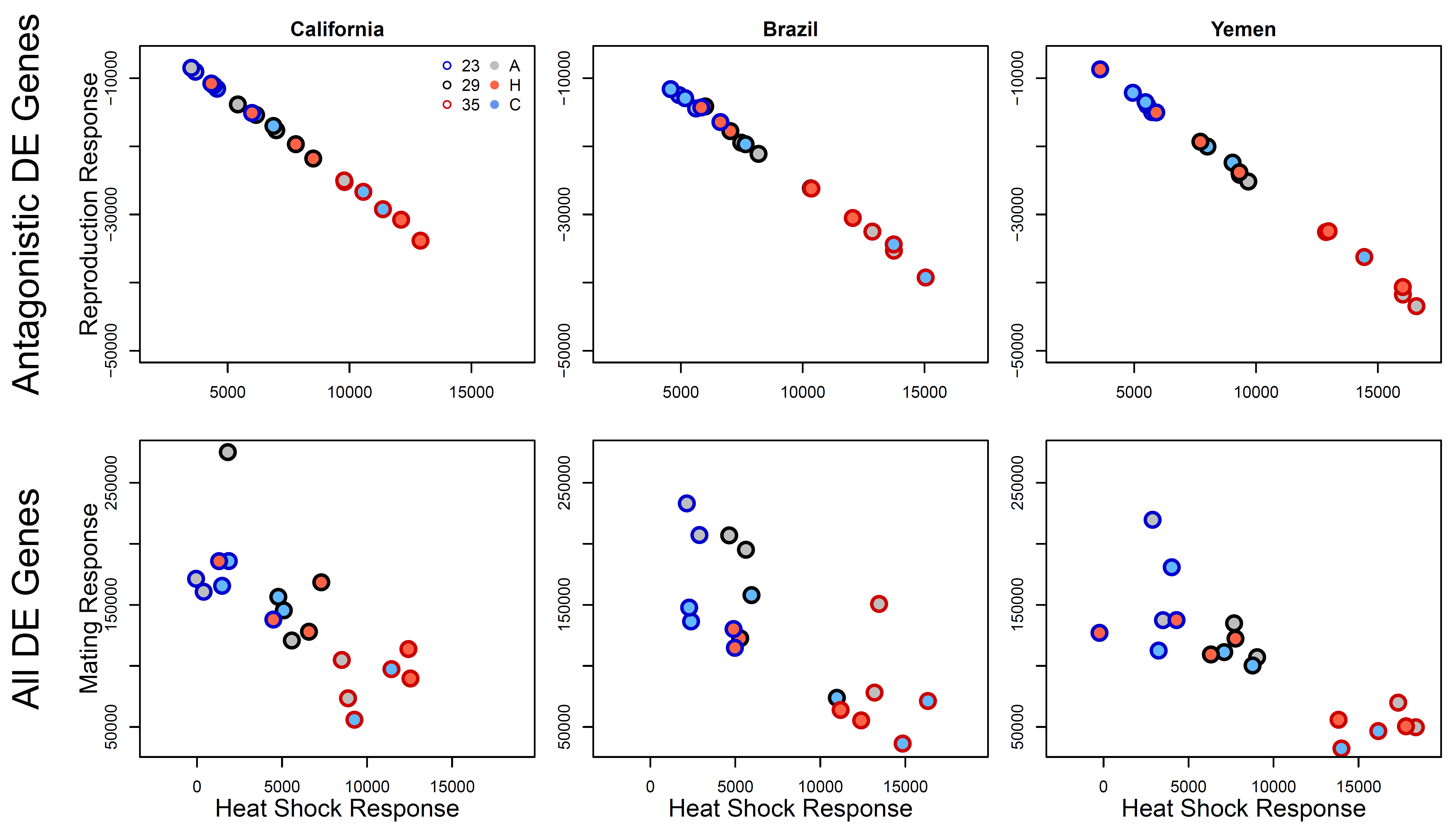

**Figure S3: Allocation trade-off between reproduction and heat stress response in experimental evolution lines based on all significant genes.**

Top panels: expression responses based on the 134 antagonistic genes in hot-adapted (red fill), cold-adapted (blue fill) and ancestors (grey fill), reared at 23 (blue border), 29 (black border) or 35°C (red border). Bottom panels: the same patterns for analyses based on all genes differentially expressed in response to heat stress (765 genes) and mating (1269 genes). There was not general effect of evolution regime across the three backgrounds in either analysis, but a strong effect of rearing temperature, with allocation towards stress response and away from reproduction at 35°C.

**Supplementary 3:**

univariate models of thermal plasticity and adaptation in life history traits

```
############ THERMAL PLASTICITY: in Ancestral founders ###################

### Lifetime Reproductive Success ###
offspring ~ temp * origin + experiment * temp + (1 | date)
Analysis of Deviance Table (Type II Wald F tests with Kenward-Roger df)

                       F Df Df.res    Pr(>F)
temp             33.5466  2  7.029 0.0002538 ***
origin            9.5749  2 98.301 0.0001588 ***
experiment       18.0062  2  7.672 0.0012656 **
temp:origin       2.5896  4 73.275 0.0435995 *
temp:experiment  14.2318  4  8.365 0.0008606 ***


### Development Time ###
dev ~ origin * temp

              Sum Sq Df    F value    Pr(>F)
origin          0.24  2    11.8464  0.003013 **
temp         1131.34  2 56919.7122 < 2.2e-16 ***
origin:temp     0.06  4     1.5213  0.275500
Residuals       0.09  9


### Metabolic Rate ###
log(co2) ~ origin * temp + log(aw.mean) + (1 | Date) + (1 | Tube)
Analysis of Deviance Table (Type II Wald F tests with Kenward-Roger df)

                    F Df  Df.res    Pr(>F)
origin         2.0253  2 102.021 0.1372207
temp         107.8867  2   3.456 0.0007681 ***
log(aw.mean)  96.3264  1  93.599 4.745e-16 ***
origin:temp    1.7845  4  93.323 0.1385070


### Adult Weight ###
aw.before ~ origin * temp2 + (1 | Date)
Analysis of Deviance Table (Type II Wald F tests with Kenward-Roger df)
                  F Df  Df.res    Pr(>F)
origin       3.8808  2 103.965 0.0236895 *
temp        28.1112  2   2.744 0.0148681 *
origin:temp  6.2839  4 103.983 0.0001437 ***


### Early Fecundity ###
eggs ~ origin * temp2 + (1 | Tube) + (1 | Date)

Analysis of Deviance Table (Type II Wald F tests with Kenward-Roger df)
                 F Df Df.res   Pr(>F)
origin      4.7965  2 90.591 0.010471 *
temp        8.2937  2  2.949 0.061544 .
origin:temp 4.1946  4 98.993 0.003522 **

### Water loss ###
log(wvp.corr) ~ origin * temp2 + (1 | Tube)
Analysis of Deviance Table (Type II Wald F tests with Kenward-Roger df)

                 F Df Df.res Pr(>F)
origin      1.3167  2 96.187 0.2728
temp      262.1866  2 78.984 <2e-16 ***
origin:temp 1.1731  4 87.864 0.3282


### Weigth Loss ###
log(aw.loss) ~ origin * temp2 + log(aw.mean) + (1 | Tube) + (1 | Date)
Analysis of Deviance Table (Type II Wald F tests with Kenward-Roger df)

                  F Df  Df.res  Pr(>F)
origin       2.5414  2  91.487 0.08431 .
temp         8.3343  2   3.792 0.04093 *
log(aw.mean) 0.0008  1 100.813 0.97774
origin:temp  2.0975  4  98.218 0.08687 .


############ ADAPTATION:  Evolution Regime Differences ###################

### Lifetime Reproductive Success ###
offspring ~ temp*origin*regime+experiment * temp + (1|pop)+(1| pop:temp)
Analysis of Deviance Table (Type II Wald F tests with Kenward-Roger df)

                             F Df  Df.res    Pr(>F)
temp                   22.9443  1    6.81 0.0021500 **
origin                  0.7091  2    5.99 0.5291548
regime                 48.9568  1    6.23 0.0003605 ***
experiment              8.7884  2 1023.46 0.0001643 ***
temp:origin             2.0129  2    6.02 0.2141043
temp:regime             1.2883  1    6.37 0.2972912
origin:regime           0.6842  2    6.00 0.5399057
temp:experiment        19.6974  2 1024.06 4.036e-09 ***
temp:origin:regime      0.4912  2    6.02 0.6344217


### Development Time ###
devtime~temp*origin*regime + (1|pop) + (1|pop:temp)
Analysis of Deviance Table (Type II Wald F tests with Kenward-Roger df)

                           F Df Df.res    Pr(>F)
temp               8926.9130  1      6 9.472e-11 ***
origin                8.4124  2      6   0.01817 *
regime               10.5487  1      6   0.01751 *
temp:origin           5.0516  2      6   0.05173 .
temp:regime         108.4421  1      6 4.595e-05 ***
origin:regime         0.2305  2      6   0.80084
temp:origin:regime    2.0679  2      6   0.20743


### Metabolic Rate ###
log(co2)~temp*origin*regime+log(aw)+ (1|pop)+(1|pop:temp)+(1|date)+(1|tube)
Analysis of Deviance Table (Type II Wald F tests with Kenward-Roger df)

                              F Df Df.res    Pr(>F)
temp                   623.9250  1  12.55 4.435e-12 ***
origin                   1.0236  2   6.36   0.41175
regime                   1.8975  1  13.67   0.19050
log(aw)                185.8547  1 328.30 < 2.2e-16 ***
temp:origin              2.1438  2   6.50   0.19270
temp:regime              1.1742  1  12.82   0.29850
origin:regime            2.1009  2   6.54   0.19735
temp:origin:regime       4.8510  2   7.16   0.04659


### Adult Weight ###
aw ~ temp2 * origin * regime + (1 | pop) + (1 | pop:temp)
Analysis of Deviance Table (Type II Wald F tests with Kenward-Roger df)

                             F Df Df.res   Pr(>F)
temp                    4.5182  1 5.6231 0.080762 .
origin                 13.2373  2 5.9595 0.006420 **
regime                 70.9266  1 5.9626 0.000158 ***
temp:origin             7.5042  2 5.8614 0.024192 *
temp:regime            11.5186  1 5.9896 0.014644 *
origin:regime           2.4900  2 6.0374 0.162713
temp:origin:regime      3.7048  2 6.1139 0.088317 .


### Early Fecundity ###
eggs~temp*origin*regime+egg.counter+(1|pop)+(1|pop:temp)+(1|date)+(1|tube)
Analysis of Deviance Table (Type II Wald F tests with Kenward-Roger df)

                             F Df Df.res    Pr(>F)
temp                   78.7184  1  17.11 8.229e-08 ***
origin                 16.9281  2   6.07  0.003293 **
regime                 11.5846  1  17.08  0.003362 **
egg.counter            35.4965  1 343.36 6.331e-09 ***
temp:origin             0.5990  2   6.22  0.578234
temp:regime             8.2913  1  17.14  0.010344 *
origin:regime           2.6100  2   7.32  0.139386
temp:origin:regime      0.5344  2   7.37  0.607089


### Water loss ###
log(wvp)~temp*origin*regime+log(aw)+(1 |pop)+(1|pop:temp)+(1|date)+(1|tube)
Analysis of Deviance Table (Type II Wald F tests with Kenward-Roger df)

                              F Df  Df.res    Pr(>F)
temp                   941.0332  1   6.947 1.123e-08 ***
origin                   1.5900  2   5.752   0.28203
regime                  10.1036  1   8.310   0.01240 *
log(aw)                  0.9647  1 289.272   0.32684
temp:origin              0.2615  2   5.592   0.77884
temp:regime              9.3107  1   7.250   0.01777 *
origin:regime            0.3662  2   5.751   0.70845
temp:origin:regime       0.8626  2   5.827   0.46975


### Weigth Loss ###
log(weight.loss+1e-04)~temp*origin*regime+log(aw.mean)+(1|pop)+(1|pop:temp)+(1|date)+(1|tube)
Analysis of Deviance Table (Type II Wald F tests with Kenward-Roger df)

                            F Df Df.res   Pr(>F)
temp                   72.6328  1  18.35 8.47e-08 ***
origin                 10.3677  2   6.31  0.01012 *
regime                  0.9370  1  19.04  0.34519
log(aw.mean)            5.5838  1 359.36  0.01866 *
temp:origin             2.3715  2   6.26  0.17110
temp:regime             7.3232  1  17.93  0.01450 *
origin:regime           0.3279  2   6.74  0.73130
temp:origin:regime      1.1515  2   6.79  0.37102
```

**Supplementary 4**:

Alternative estimates of laboratory fitness in experimental evolution lines

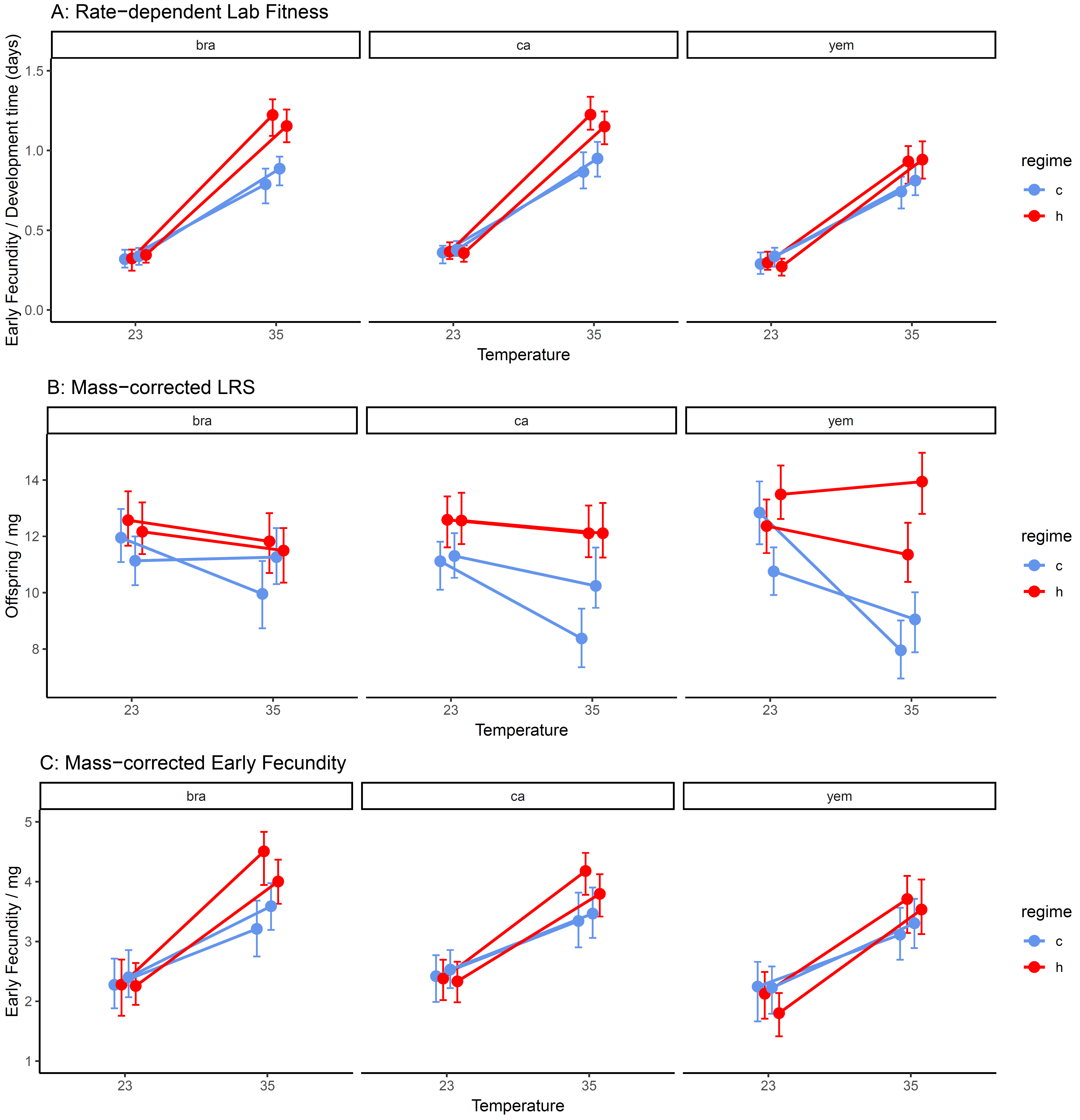

**Figure S4**: In **A)** as a proxy for fitness of each line at the conditions and two thermal conditions used in experimental evolution, we calculated a rate-dependent measure as early fecundity per female divided by development time (which corresponds roughly to generation time; mean time from start of egg laying until the emergence of reproductively mature adults) from Dataset 2. As eggs laid early should have an advantage due to being first to utilize the host resource, the calculation should give a reasonable estimate of laboratory fitness during experimental evolution. In **B**) LRS is shown corrected for mean body mass of females and in **C**) early fecundity corrected for mass is shown. All three measures show that hot-adapted lines outcompete cold-adapted lines at 35°C. Differences are less clear at 23°C, even after correcting for the larger size of females from hot-adapted lines. Whiskers represent 95% CIs based on parametric bootstrap.

**Supplementary 5**:

Summary of models of host consumption in experimental evolution lines

```
####### Analyses of host consumption in hot and cold regime ############

### Total host biomass consumed:
loss.ctrl.fem ~ regime*origin*temp + (1|pop) + (1|temp:pop)
Analysis of Deviance Table (Type II Wald F tests with Kenward-Roger df)

                             F Df  Df.res   Pr(>F)
regime                 22.8112  1  5.9715 0.003115 **
origin                  0.5881  2  5.9551 0.584685
temp                    0.5359  2 11.7887 0.598759
regime:origin           1.3612  2  5.9962 0.325530
regime:temp             3.2465  2 11.8820 0.075016 .
origin:temp             0.8559  4 11.8547 0.517510
regime:origin:temp      0.4674  4 11.9324 0.758747


### Consumed host biomass per beetle emerging:
cons.beetle ~ regime*origin*temp + (1|pop) + (1|temp2:pop)
Analysis of Deviance Table (Type II Wald F tests with Kenward-Roger df)

                             F Df  Df.res   Pr(>F)
regime                 16.3913  1  5.9410 0.006877 **
origin                  0.6981  2  5.9144 0.534344
temp                    1.9469  2 11.5953 0.186625
regime:origin           1.4082  2  5.9818 0.315383
regime:temp             0.5502  2 11.7669 0.590976
origin:temp             0.6558  4 11.7140 0.634330
regime:origin:temp      0.6467  4 11.8705 0.639931


### Consumed host biomass per egg laid:
cons.egg ~ regime*origin*temp + (1|pop) + (1|temp:pop)
Analysis of Deviance Table (Type II Wald F tests with Kenward-Roger df)

                             F Df  Df.res    Pr(>F)
regime                 15.2788  1  5.9118  0.008137 **
origin                  0.0864  2  5.8858  0.918403
temp                   34.6239  2 11.4288 1.412e-05 ***
regime:origin           0.4961  2  5.9513  0.632026
regime:temp             0.0631  2 11.6719  0.939182
origin:temp             0.9797  4 11.5938  0.455681
regime:origin:temp      3.3172  4 11.8327  0.048229 *


### Consumed host biomass per beetle biomass:
cons.biomass ~ regime*origin*temp + (1|pop) + (1|temp:pop)
Analysis of Deviance Table (Type II Wald F tests with Kenward-Roger df)

                             F Df  Df.res   Pr(>F)
regime                  0.5985  1  5.7284 0.469870
origin                  1.3658  2  5.6298 0.328431
temp                   10.5747  2 10.7792 0.002873 **
regime:origin           0.6121  2  5.7766 0.573975
regime:temp             2.6759  2 11.0591 0.112765
origin:temp             0.4299  4 10.8771 0.784332
regime:origin:temp      1.0298  4 11.1742 0.433849
```

```
####### Analyses of beetle traits in hot and cold regime ############

### Minimum development time of emerging beetles:
NB.DAYS.DVLP ~ regime*origin*temp + (1|pop) + (1|temp:pop)
Analysis of Deviance Table (Type II Wald F tests with Kenward-Roger df)

                              F Df Df.res    Pr(>F)
regime                   0.9963  1      6   0.35675
origin                   1.1980  2      6   0.36494
temp                   754.7321  1      6 1.538e-07 ***
regime:origin            3.5293  2      6   0.09700 .
regime:temp              1.4842  1      6   0.26885
origin:temp2             1.3395  2      6   0.33040
regime:origin:temp       3.9220  2      6   0.08141 .


### Mean body mass of emerging beetles:
beetle.size ~ regime*origin*temp + (1|pop) + (1|temp:pop)
Analysis of Deviance Table (Type II Wald F tests with Kenward-Roger df)

                              F Df Df.res    Pr(>F)
regime                  21.8705  1 5.8145 0.0036987 **
origin                   2.6164  2 5.7760 0.1552477
temp                   131.3228  1 4.7673 0.0001191 ***
regime:origin            0.5676  2 5.8428 0.5952821
regime:temp              7.8662  1 5.0982 0.0369797 *
origin:temp             12.4183  2 4.7663 0.0128767 *
regime:origin:temp       1.1448  2 5.1029 0.3884123
```

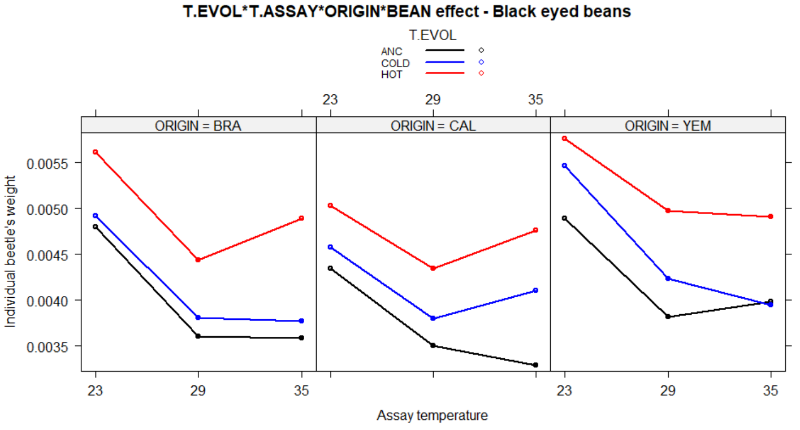

*Note that the ancestors (black) are included for illustrative purposes but were not included in the statistical analyses presented in the anova tables*

**Figure S5:** Mean body mass of beetles emerging from the assays of host consumption.

```
### Model explaining host consumption from life history traits:
#fecundity ="eggs"" #Mean adult mass ="beetle.size" #juvenile survival = "survival"
loss.ctrl ~ scale(beetle.size)+scale(survival)+scale(eggs)+temp

                   Estimate Std. Error t value Pr(>|t|)
(Intercept)         0.97480    0.01037  94.027  < 2e-16 ***
scale(beetle.size)  0.07982    0.01121   7.117 2.08e-10 ***
scale(survival)     0.10423    0.01117   9.331 4.45e-15 ***
scale(eggs)         0.41430    0.01309  31.641  < 2e-16 ***

Residual standard error: 0.1031 on 95 degrees of freedom
  (10 observations deleted due to missingness)
Multiple R-squared:  0.9273,	Adjusted R-squared:  0.925
F-statistic: 403.7 on 3 and 95 DF,  p-value: < 2.2e-16
```

**Supplementary 6**:

Selection on female body mass at different temperatures

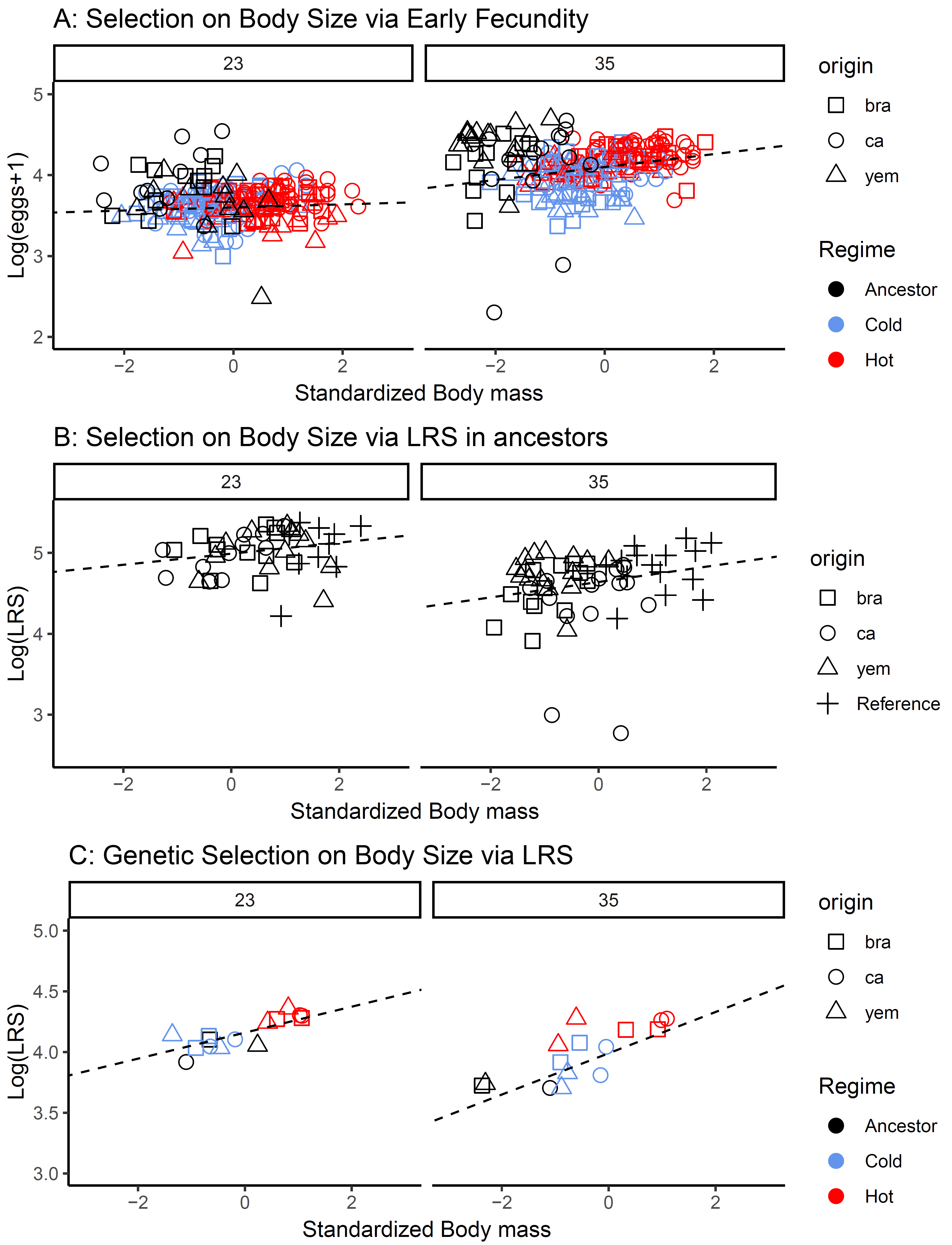

**Figure S6:** **A**) Regression of logarithmized early fecundity on variance standardized body mass for each female trio from dataset 1 & 2. **B**) Regression of logarithmized lifetime reproductive success (number of adult offspring produced) and variance standardized body mass for female trios from the experiment on ancestral populations that measured both traits on the same individuals. **C**) Regression of logarithmized mean lifetime reproductive success on mean (variance standardized) body mass for each line used in the experiment. Selection gradients on size are given by the regression slopes. These slopes were estimated across all data. In panel A, this was done after first removing effects of evolution regime (cold/hot/ancestor). In each case the selection gradient is steeper at 35°C compared to 23°C.

```
### Selection on body mass (aw) via early fecundity in evolution lines ###
### Corresponds to panel A of Figure S6 ###


log(eggs + 1) ~ temp*scale(aw)+selection*temp

                         Estimate Std. Error t value Pr(>|t|)
(Intercept - ancestor)   3.82066    0.04461  85.636  < 2e-16 ***
temp35                   0.47389    0.06821   6.948 1.29e-11 ***
scale(aw)                0.02080    0.02303   0.903  0.36694
regime.c                -0.20117    0.04993  -4.029 6.56e-05 ***
regime.h                -0.18576    0.05943  -3.126  0.00189 **
temp35:scale(aw)         0.05512    0.03186   1.730  0.08432 .
temp35:regime.c         -0.13462    0.07497  -1.796  0.07322 .
temp35:regime.h          0.04767    0.08700   0.548  0.58399

Residual standard error: 0.2475 on 456 degrees of freedom
Multiple R-squared:  0.496,	Adjusted R-squared:  0.4882
F-statistic:  64.1 on 7 and 456 DF,  p-value: < 2.2e-16


Anova Table (Type II tests)
                 Sum Sq  Df  F value    Pr(>F)
temp            21.7727   1 355.4800 < 2.2e-16 ***
scale(aw)        0.5945   1   9.7067  0.001952 **
regime           3.6228   2  29.5747  8.38e-13 ***
temp:scale(aw)   0.1833   1   2.9927  0.084316 .
temp:regime      0.7136   2   5.8252  0.003176 **
Residuals       27.9294 456


#at 23C:

            Estimate Std. Error t value Pr(>|t|)
(Intercept)  3.82377    0.04125  92.707  < 2e-16 ***
scale(aw)    0.01923    0.01912   1.006 0.315681
reimge.c    -0.20117    0.04484  -4.486 1.15e-05 ***
regime.h    -0.18576    0.05337  -3.480 0.000601 ***

Residual standard error: 0.2223 on 226 degrees of freedom
Multiple R-squared:  0.08268,	Adjusted R-squared:  0.0705
F-statistic:  6.79 on 3 and 226 DF,  p-value: 0.0002109


#at 35C:

            Estimate Std. Error t value Pr(>|t|)
(Intercept)  4.28339    0.05393  79.420  < 2e-16 ***
scale(aw)    0.07973    0.02523   3.160  0.00179 **
regime.c    -0.33578    0.06101  -5.504  9.9e-08 ***
regime.h    -0.13808    0.06932  -1.992  0.04755 *

Residual standard error: 0.27 on 230 degrees of freedom
Multiple R-squared:  0.1872,	Adjusted R-squared:  0.1765
F-statistic: 17.65 on 3 and 230 DF,  p-value: 2.396e-10


### Selection on body mass (aw) via LRS in ancestors ###
### Corresponds to panel B of Figure S6 ###


log(total.fecundity+1) ~ temp *scale(aw.before)

                         Estimate Std. Error t value Pr(>|t|)
(Intercept)               4.98807    0.06024  82.809  < 2e-16 ***
temp35                   -0.34478    0.07855  -4.389 2.85e-05 ***
scale(aw.before)          0.06846    0.06275   1.091    0.278
temp35:scale(aw.before)   0.02664    0.07984   0.334    0.739

Residual standard error: 0.3645 on 99 degrees of freedom
Multiple R-squared:  0.2666,	Adjusted R-squared:  0.2443
F-statistic: 11.99 on 3 and 99 DF,  p-value: 9.177e-07


Anova Table (Type II tests)

Response: log(total.fecundity + 1)
                        Sum Sq Df F value    Pr(>F)
temp                    2.5541  1 19.2210 2.905e-05 ***
scale(aw.before)        0.6367  1  4.7919   0.03094 *
temp:scale(aw.before)   0.0148  1  0.1114   0.73928
Residuals              13.1551 99


#at 23C:

                 Estimate Std. Error t value Pr(>|t|)
(Intercept)       5.01642    0.03976 126.179   <2e-16 ***
scale(aw.before)  0.05996    0.04021   1.491    0.143

Residual standard error: 0.2667 on 43 degrees of freedom
Multiple R-squared:  0.04917,	Adjusted R-squared:  0.02706
F-statistic: 2.224 on 1 and 43 DF,  p-value: 0.1432


#at 35C:

                 Estimate Std. Error t value Pr(>|t|)
(Intercept)       4.61273    0.05575  82.733   <2e-16 ***
scale(aw.before)  0.09304    0.05624   1.654    0.104

Residual standard error: 0.4246 on 56 degrees of freedom
Multiple R-squared:  0.04659,	Adjusted R-squared:  0.02956
F-statistic: 2.736 on 1 and 56 DF,  p-value: 0.1037
```

```
### Genetic selection on body mass (aw) via LRS ###
### Corresponds to panel C of Figure S6 ###


log(LRS) ~ temp*scale(aw)

                  Estimate Std. Error t value Pr(>|t|)
(Intercept)        4.12760    0.03152 130.946  < 2e-16 ***
temp35           -0.10609    0.04432  -2.394  0.02418 *
scale(aw)          0.12201    0.03590   3.398  0.00219 **
temp235:scale(aw)  0.03371    0.04608   0.731  0.47107

Residual standard error: 0.118 on 26 degrees of freedom
Multiple R-squared:  0.6828,	Adjusted R-squared:  0.6462
F-statistic: 18.65 on 3 and 26 DF,  p-value: 1.158e-06


Anova Table (Type II tests)

Response: log(LRS)
                 Sum Sq Df F value    Pr(>F)
temp            0.07752  1   5.570   0.02605 *
scale(aw)       0.55746  1  40.057 1.059e-06 ***
temp:scale(aw)  0.00744  1   0.535   0.47107
Residuals       0.36183 26


#At 23C:

            Estimate Std. Error t value Pr(>|t|)
(Intercept)  4.15518    0.01961  211.93  < 2e-16 ***
scale(aw)    0.10715    0.02029    5.28 0.000149 ***

Residual standard error: 0.07593 on 13 degrees of freedom
Multiple R-squared:  0.682,	Adjusted R-squared:  0.6575
F-statistic: 27.88 on 1 and 13 DF,  p-value: 0.000149


#At 35C:

            Estimate Std. Error t value Pr(>|t|)
(Intercept)  3.98632    0.03836  103.93  < 2e-16 ***
scale(aw)    0.16991    0.03970    4.28 0.000896 ***

Residual standard error: 0.1485 on 13 degrees of freedom
Multiple R-squared:  0.5849,	Adjusted R-squared:  0.5529
F-statistic: 18.32 on 1 and 13 DF,  p-value: 0.0008964
```
